# MicroPET evidence for a hypersensitive neuroinflammatory profile of gp120 mouse model of HIV

**DOI:** 10.1101/2021.10.11.463989

**Authors:** Jared W. Young, Christopher V. Barback, Louise A. Stolz, Stephanie M. Groman, David R. Vera, Carl Hoh, Kishore K. Kotta, Arpi Minassian, Susan B. Powell, Arthur L. Brody

## Abstract

Despite increased survivability for people living with HIV (PLWH), HIV-related cognitive and behavioral abnormalities persist. Determining the biological mechanism(s) underlying these abnormalities is critical to minimize the long-term impact of HIV. Human positron emission tomography (PET) studies reveal that PLWH exhibit higher neuroinflammation, which may contribute to cognitive and behavioral problems. PLWH are hypersensitive to environmental insults that drive elevated inflammatory profiles. Gp120 is an envelope glycoprotein exposed on the surface of the HIV envelope which enables HIV virus entry into a cell and contributes to HIV-related neurotoxicity. Gp120 overexpression in mice enables delineating its impact, including on neuroinflammation. I*n vivo* evidence for gp120 transgenic (Tg) mice exhibiting neuroinflammation has yet to be determined.

Here, we conducted microPET imaging in male gp120 Tg and wildtype mice, using the radiotracer [(18)F]FEPPA which binds to the translocator protein expressed by activated microglial and serves as a marker of neuroinflammation. Imaging was performed at baseline and 24 hours after treatment with lipopolysaccharide (LPS; 5 mg/kg), endotoxin that triggers an immune response.

Gp120 Tg mice exhibited elevated [(18F)]FEPPA in response to LPS vs. wildtype mice throughout the brain including dorsal and ventral striata, hypothalamus, and hippocampus, but not prefrontal cortex.

Gp120 Tg mice are hypersensitive to environmental inflammatory insults, consistent with PLWH, measurable *in vivo*. It remains to-be-determined whether this heightened sensitivity is connected to the behavioral abnormalities of these mice or is sensitive to antiretroviral or other treatments.

## Introduction

Human Immunodeficiency Virus (HIV) has transitioned from a uniformly fatal illness to a chronic – albeit still life-threatening – disease, with the development of antiretroviral therapy (ART). People living with HIV (PLWH), still suffer from cognitive and behavioral abnormalities that negatively impact their lives and is referred to as HIV-associated neurocognitive disorders (HAND) ^1-3^. PLWH also exhibit hypersensitivity to environmental insults, e.g., infectious agents and medications (prophylactics, antibiotics, drugs to treat infections, and antiretrovirals ^4^). The mechanisms driving these changes is not known, but cognitive deficits have been linked to neuroinflammation in several diseases associated with neural insults ^5, 6^, including HIV ^7-10^.

Even in people with near-zero virologic expression, inflammatory markers are elevated in the peripheral blood, such as IL-6 ^11^, and in the brain, as measured by positron emission tomography (PET) and the translocator protein (TSPO; a marker of microglial activation) ^12^. Despite ART-induced increase in life expectancies, prevalence of cognitive deficits remain ^13^ and may be linked to persistent neuroinflammation^14, 15^. HIV can enter the brain during early stages of infection through infected lymphocytes and monocytes, and neurons can be further affected through loss of synapses and cell death via pro-inflammatory cytokines ^2^. Therefore it remains critical to determine the mechanism(s) underlying this neuroinflammation to minimize the impact of HIV on the lives of PLWH.

Neuroinflammation occurs in the brain and is observable *in vivo* PET ^8, 12, 16^ and *ex vivo* postmortem studies^17^ of PLWH. Such neuroinflammation may be critical to cognitive and behavioral function in PLWH, as it may correlate to such abnormalities in PLWH ^12^. PET radiotracers enable the examination of a marker for neuroinflammation by labeling activated microglia *in vivo* ^18^. Microglia are a central component of the neuroinflammatory response, and, when activated, express the TSPO 18 kDa, thereby making levels of this molecule a primary marker for neuroinflammation. TSPO is a mitochondrial protein found in activated microglia and can be used to study patterns of neuroinflammation ^19, 20^. Studies using TSPO PET tracers have found higher TSPO binding in the thalamus, frontal cortex, temporal and occipital cortex, and the putamen in individuals with HAND compared to healthy controls ^21^. [(18)F]FEPPA is one such PET tracer and has been demonstrated to reliably bind to TSPO ^22-26^. [(18)F]FEPPA is available for studies in humans and rodents ^22, 24^, and so enables translational research directed at understanding the biological mechanisms driving this neuroinflammatory profile relevant to HIV.

Gp120 is the glycoprotein that enables virus entry into a cell contributing to HIV-related neurotoxicity and its overexpression enables determining the impact of elevated levels as seen in HIV. Gp120 transgenic (Tg) mice exhibit several HIV-relevant features, including increased *in vitro* neuroinflammatory measures of GFAP, Iba-1, IL6 ^27, 28^, as well as behavioral, cognitive and neurophysiological deficits ^29, 30^, including impaired reinforcement learning, executive functioning,^31^ risk-taking, and drug sensitivity^30^. To date, confirmation of gp120-induced *in vivo* neuroinflammation in these mice has yet to be determined. Confirmation of such a neuroinflammatory profile in gp120 mice – especially a hypersensitivity to environmental challenge such as infections – would: 1) confirm the importance of gp120 in HIV-relevant neuroinflammation; 2) enable longitudinal studies determining the interactive drug abuse effect on such neuroinflammation and subsequent HIV-relevant behaviors (given elevated substance abuse in PLWH); and 3) identify the gp120 Tg mice as a model system by which to develop therapeutic targets to limit this neuroinflammatory profile. Here, we assessed neuroinflammation via TSPO quantification with [(18)F]FEPPA radiotracer and microPET imaging in male gp120 Tg and wildtype mice before and after administration of the inflammatory agent lipopolysaccharide (LPS; 5 mg/kg). LPS was chosen because it activates Toll-like receptor 4 primarily expressed in microglia resulting in the production of pro-inflammatory cytokines and TSPO-measured neuroinflammation ^23, 32, 33^.

## Materials and Methods

### Animals

Gp120 expression in these Tg mice occur in astrocytes under the control of a modified murine glial fibrillary acidic protein (GFAP) promoter. These mice were generated from a mixed C57BL/6 x Sv129 (SJL/BL6/129) background ^34, 35^. For the current study, Tg mice on a mixed BL6/129 background were crossed with BDF1 mice (both male and female) obtained from Charles River. Non-Tg littermates (wildtype, WT) were used as controls. Male mice from 8–12 months-old Tg and WT littermates (n=7 and 6 per condition respectively, from an F10 BL6/BDF1 generation) were used for the current study, weighing 27-37 g at the beginning of the study. Confirmation of genotype was conducted by Transnetyx via PCR analysis of tail DNA.

Mice were housed in groups of 1–4 per cage in a climate-controlled animal colony on a reversed day/night cycle (lights on at 8:00 PM, off at 8:00 AM). For microPET studies, mice were transferred to the room containing the equipment 24 h before testing began (see Fig. 1), with free access to food (Haran Teklad, Madison, WI) and water throughout. All testing was conducted between 1:00 and 6:00 PM. All procedures were approved by the UCSD Institutional Animal Care and Use Committee and conformed to NIH guidelines.

**Figure 1.**
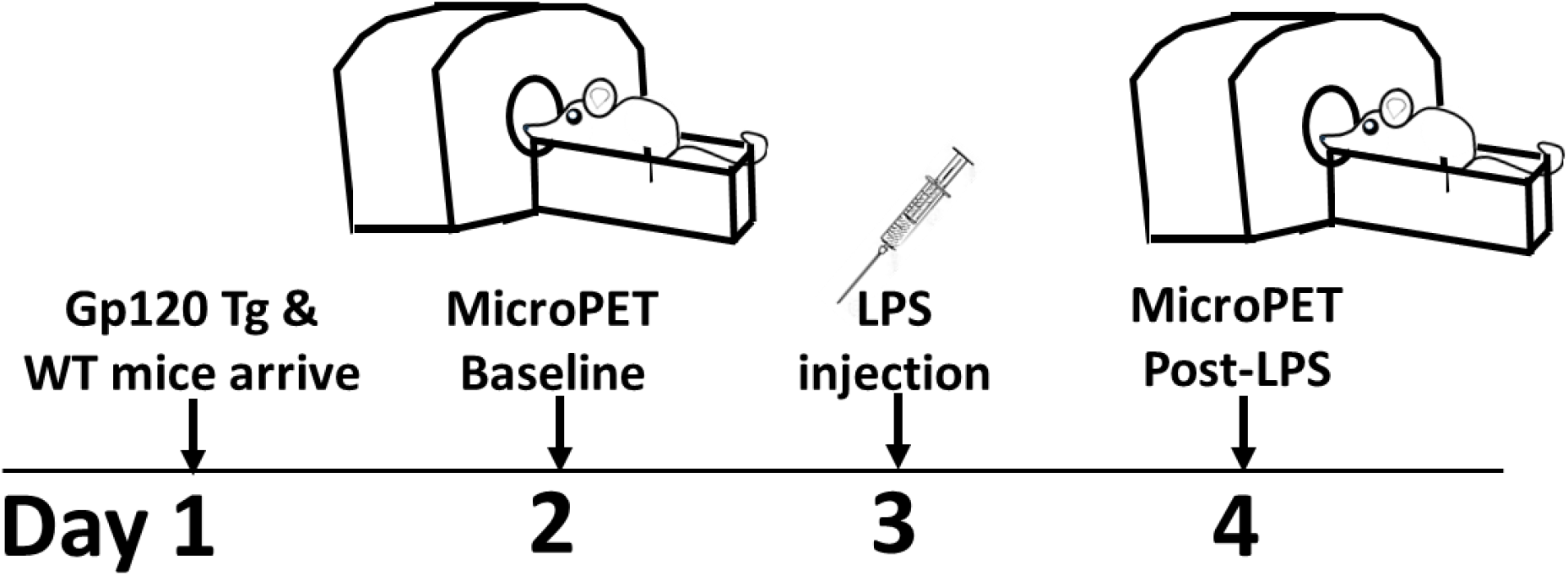
gp120 microPET study timeline: Gp120 transgenic (Tg) and wildtype (WT) littermates arrive at the Micro-PET center on Day 1. After 24 h of recovery from travel, mice receive a single bolus injection of ^18^F-labeled FEPPA via the tail vein, carrying between 100 and 300 μCi of activity in a volume of 500-1000 μL, and scanned on a microPET to quantify baseline levels of TSPO. 24 h later, mice are injected with lipopolysaccharide (LPS; 5 mg/kg). The mice are then re-scanned 24 h later to quantify LPS-induced changes in TSPO (Day 4). At the conclusion of each micro-PET scan, mice were moved to the bed of a micro-MRI scanner, where T1 and T2 weighted images of the head were acquired along all three axes.

### Reagents

FEPPA precursor and FEPPA standards were purchased from Huyai Isotopes (city and country). At the time of arrival analytical HPLC (Phenomenex Luna 5 μm C18(2) 100 A° 250 × 4.6mm) was performed on FEPPA precursor to measure the chemical purity using a mobile phase of 80:20 Methanol:Water, 0.1% Formic acid, flow rate (1.0 mL/min). The infectious agent chosen was Lipopolysaccharide (LPS, 5 mg/kg; Escherichia coli O111:B4; Sigma Aldrich, MO, USA), because of our previous experience with this agent ^36^, and the dose chosen induced elevated neuroinflammation in rats ^20^.

### [(18)F]FEPPA Synthesis

Synthesis of [(18)F]FEPPA was performed according to the previously^37^ published paper using GE FXN Radiochemistry synthesis unit. The [^18^F] fluoride was transferred from the target to the synthesis module (FXN) using inert gas pressure. The [^18^F] was separated from the [^18^O]-water by trapping the [^18^F] on an anion exchange cartridge (QMA carbonate light cartridge). The [^18^F]-ions were then eluted from the QMA cartridge into the labeling vessel using a solution containing Kryptofix/K_2_CO_3_ in methanol/acetonitrile mixture. The solvent was then evaporated. The tosyl precursor in acetonitrile was added to the dry residue and a nucleophilic substitution reaction was run under 120 ^º^C and then under 60 °C to obtain crude [(18)F]FEPPA, which was subsequently purified by preparative HPLC (Phenomenex Luna 5 μm C18(2) 100 A° 250 × 10m
m) using 50% methanol and 50% sterile water containing 0.5% (v/v) formic acid as an eluent (flow rate 3 ml/min) to obtain [(18)F]FEPPA with >98% radiochemical purity and >5 Ci/μmole specific activity.

### Experimental Micro-PET imaging Design

On the first day of imaging, each mouse was anesthetized with isoflurane, weighed (as an estimate of volume), and placed upon the micro-PET (Vista DR from GE Healthcare) gantry bed. They were advanced into the imager until their heads rested in the center of the field of view. The microPET scanner was set to take a single dynamic scan of each mouse, over 20 min. The scan began just before the imaging agent was injected. Once the scan commenced, each mouse received a single bolus injection of ^18^F labeled FEPPA via the tail vein, carrying between 100 and 300 μCi of activity in a volume of 500-1000 μL. [(18)F]FEPPA was chosen as the radiotracer for this study because its relatively long half-life (compared to 11C-labelled compounds) allows for multiple scans to be performed in the same day (and has been shown to be sensitive to LPS treatment in mice ^22^). The activity present in each syringe was measured before and after injection to provide a value for the total injected into each mouse. At the conclusion of each mouse’s microPET scan, they were moved to the bed of a microMRI scanner (4000MRS from MRSolutions), where T1 and T2 weighted images of the head were taken along all three axes. Upon completion of this scan, each mouse was allowed to recover, and the activity allowed to decay away. After 24 h, mice were injected with lipopolysaccharide (LPS; 5 mg/kg), a dose shown to increase neuroinflammation in rats ^20^. The mice were then reassessed in the microPET as described above 24 h later. Following the final microPET scan on each day, a 60mL syringe containing known activity of [(18)F]FEPPA in PBS was scanned using the same methods used in the mice. Scanning took place over 2 cohorts of animals, with 82.4uCi used for the first cohort, and 57.0uCi used for the second.

Scans were reconstructed through filtered back-projection (FBP). The FORE parameters used were a span of 3 and a Dmax of 16, and the 2D FBP used a ramp filer, with an alpha of 1.0 and a cutoff of 1.0. Random and scatter corrections were applied, but no attenuation correction was employed. Decay correction was employed throughout reconstruction. Following the reconstruction of the scans, images were registered and analyzed using a combination of ImageJ and FSL. Transaxial T2 weighted MRI images were first skull-stripped, then aligned and registered together. They were then averaged together to create an anatomical template to which PET scans and an anatomical map could also be registered. Regions of interest were constructed from this template to cover the desired brain regions. PET scans were similarly aligned, registered, and averaged to create another template for use in registering the PET scans to the MRI and anatomical mapping. Once all scans were properly registered, intensity values for each ROI at each minute of scanning were measured and compiled. Measurements of the intensity values of the 60mL syringe allowed us to equate between intensity values in the image to μCi of [(18)F]FEPPA/unit volume. Standard Uptake Values (SUVs) were calculated from these data using the following equation:

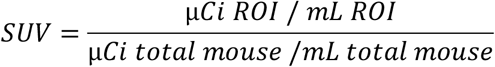

### Statistical analyses

All statistical analyses were performed using IBM SPSS Statistics 27 (Armonk, NY, USA). The SUV was the dependent variable for overall levels and for each brain region, over the whole 20 min period. Thus, data were analyzed using mixed analysis of variance (ANOVA), with pre- and post-LPS treatment and genotype as a between-subjects factors. Data were analyzed with Sphericity Assumed, with *post hoc* analyses focusing on each main or interactive effects. Results are expressed as mean ± standard error of the mean (SEM). Differences were considered statistically significant at *p*<0.05, although trends (p<0.1), were reported.

## Results

### LPS elevated neuroinflammation in gp120 Tg but not WT littermate mice, despite affecting weight equally in both genotypes

Our group used microPET and the radiotracer [(18)F]FEPPA to examine microglial activation in gp120 Tg and WT littermate mice in order to determine the *in vivo* neuroinflammatory (via elevated TSPO) profile of these mice. To confirm our ability to conduct longitudinal testing in response to drug treatment in the gp120 Tg and WT littermate mice, we assessed their neuroinflammatory profile at: 1) baseline without treatment, and 2) 24 h after administration of the pro-inflammatory agent LPS. Confirmation of the efficacy of LPS was observed given that the mice lost significant weight from baseline to testing 24 h after LPS (F(1,11)=100.0, *p*<0.0001; Fig. 2A). There was no effect of gene, or gene * LPS interaction on weight (Fs<1, ns) supporting the biological efficacy of the infectious agent in both gp120 Tg and WT mice.

**Figure 2:**
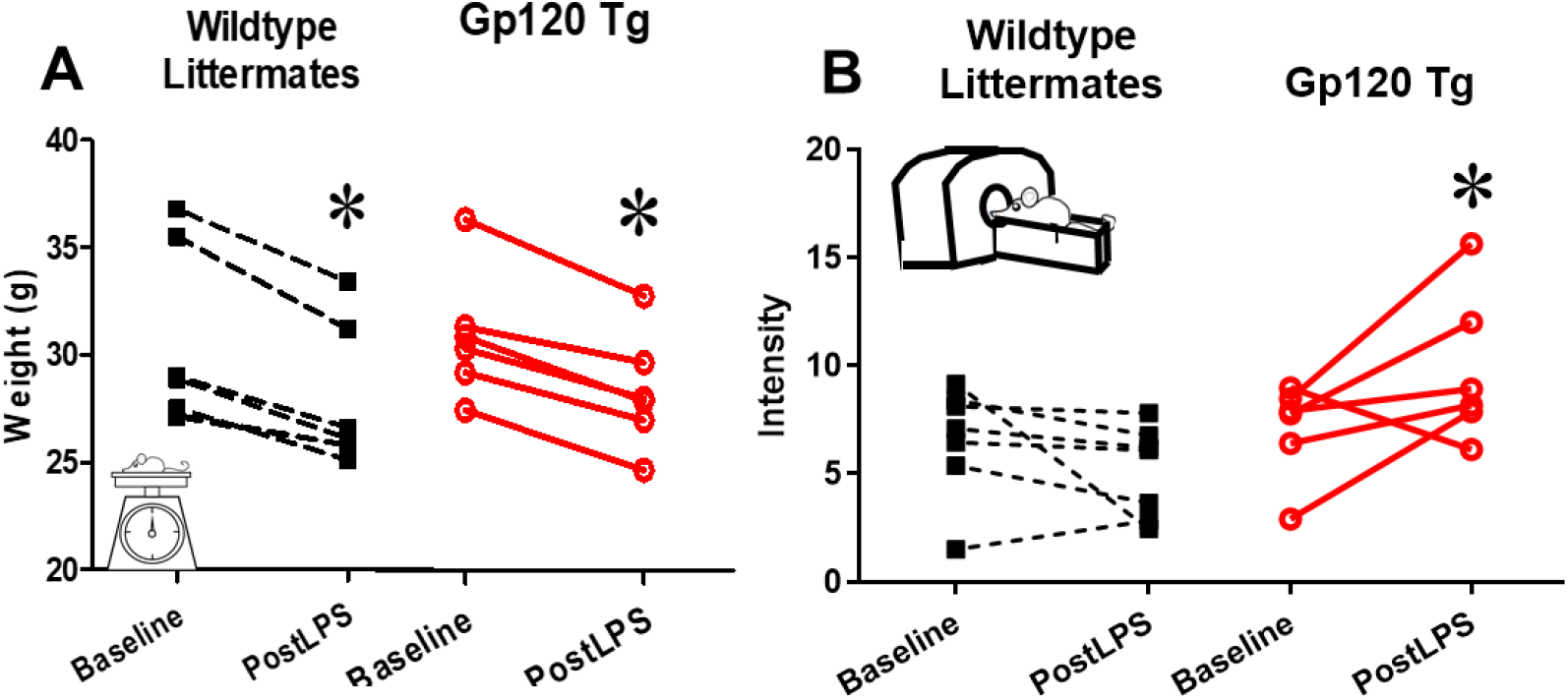
LPS induced neuroinflammation in gp120 Tg but not WT littermate mice, while equally affecting health as measured by loss in bodyweight. Gp120 transgenic (Tg; red circle and lines) and Wildtype (WT; black squares and lines) littermates were assessed at baseline and after lipopolysaccharide (PostLPS; 5 mg/kg) treatment. Importantly, LPS exerted the same level of weight loss in both Tg and WT mice (**A**). Overall at baseline, gp120 Tg mice exhibited higher microglial activation, though not significantly, while LPS increased activation specifically in gp120 Tg mice (**B**). Data presented as individual data points and their difference upon retesting. * = p<0.05.vs. baseline.

We then examined the impact of genotype and LPS on TSPO (or [(18)F]FEPPA) binding. The main effect of LPS on whole-brain SUV levels was not significant [F(1,11)=0.6, *p*=0.472], but the gene by LPS interaction was [F(1,11)=6.2, *p*<0.05; Fig. 2B]. *Post hoc* analyses revealed that LPS did not affect whole-brain SUV levels in WT mice [F(1,6)=2.4, *p*=0.175] even though these same mice lost a significant amount of weight. In gp120 mice, however, there was a trend-level increase in whole-brain SUV levels following LPS administration [F(1,5)=3.7, *p*<0.1]. Although [(18)F]FEPPA binding was higher in gp120 vs. WT mice at baseline, this effect was not significant here (F<1, ns); Fig 2), but was so 24 h following LPS injection (F_(1,11)_ =9.1, *p*<0.05) suggesting that the LPS-induced increase in [(18)F]FEPPA binding was greatest in gp120 Tg mice when compared to WT mice.

### Specific Regional Analyses

Regional differences were then examined (Fig 3) to investigate the impact of gene and LPS in a region-specific manner ^38^. A main effect of gene was observed for the dorsal striatum (F(1,11)=6.0, *p*<0.05; Fig. 3A), as well as a gene by LPS interaction (F(1,11)=5.1, *p*<0.05). *Post hoc* analyses revealed that while WT and gp120 mice did not differ at baseline (Fs<1.5, ns), gp120 mice exhibited higher SUVs in the dorsal striatum after LPS vs. WT littermate mice (*p*<0.05). Similar results were seen for the ventral striatum, with a trend of elevated SUV in gp120 tg vs. WT littermate mice [F(1,11)=3.5, *p*<0.1; Fig. 3B], with a gene by LPS interaction observed [F(1,11)=5.7, *p*<0.05]. *Post hoc* analyses revealed no effect of LPS was seen in WT mice (Fs<2, ns), nor in gp120 mice [F(1,5)=3.4, *p*=0.126], and that ventral striatal SUV levels of WT and gp120 mice did not differ at baseline (Fs<1.5, ns), gp120 mice exhibited higher SUVs than WT littermate mice after LPS treatment (*p*<0.05). Consistent with the ventral striatum, trend elevated levels of SUV were seen in the hypothalamus of gp120 vs. WT littermate mice [F(1,11)=3.5, *p*<0.1; Fig. 3C], with a gene by LPS interaction observed [F(1,11)=5.7, *p*<0.05]. Again, *post hoc* analyses revealed no LPS effect in the hypothalamus of WT (F<1, ns), or gp120 mice [F(1,5)=3.4, *p*=0.123], and while hypothalamic SUV levels of WT and gp120 mice did not differ at baseline (Fs<1.5, ns), gp120 mice exhibited higher SUVs than WT littermate mice after LPS treatment (*p*<0.05). As seen in other regions, elevated VTA SUV levels were seen in gp120 vs. WT littermate mice [F(1,11)=5.5, *p*<0.05; Fig. 3D], with a gene by LPS interaction observed [F(1,11)=5.8, *p*<0.05]. Again, *post hoc* analyses revealed no LPS effect in the VTA of WT mice (F<1.9, ns), while a LPS effect was seen in gp120 mice [F(1,5)=1.9, *p*<0.05]. While WT and gp120 mice did not differ at baseline, gp120 mice exhibited higher SUVs in the VTA (*p*<0.05). Finally, consistent with other regions, elevated PFC SUV levels were seen in gp120 vs. WT littermate mice [F(1,11)=5.6, *p*<0.05; Fig. 3E], although unlike the other brain regions, no gene by LPS interaction was observed [F(1,11)=3.0, *p*=0.112].

**Figure 3:**
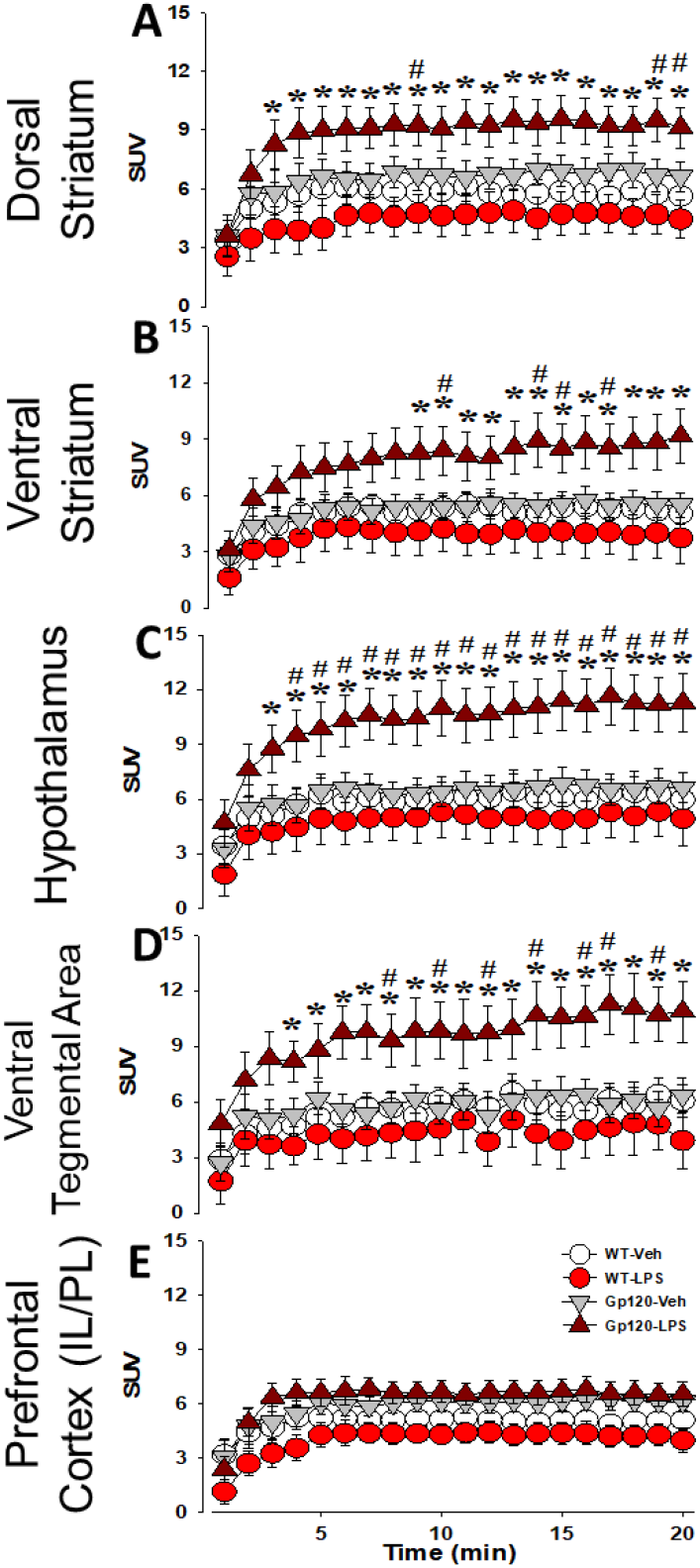
Timecourse of LPS-induced neuroinflammation in gp120 Tg mice. Elevated [(18)F]FEPPA SUV was observed in gp120 mice in response to lipopolysaccharide (LPS) across multiple brain regions, an effect not seen in their wildtype (WT) littermates. LPS-induced elevation across the 20 min imaging period was seen in the dorsal striatum (**A**), driving gp120-induced elevated levels overall in this brain region. Similar observations were made for the ventral striatum (**B**), hypothalamus (**C**), and ventral tegmental area (**D**). LPS-induced elevated [(18)F]FEPPA SUV was not seen in the prefrontal cortices of WT or gp120 Tg mice (**E**), but overall elevated levels were seen in gp120 Tg vs. WT mice. Data presented as mean ± S.E.M., * denotes *p*<0.05 vs. WT LPS-treated mice (timecourse), or vs. WT overall as indicated. # denotes *p*<0.1 vs. gp120 Veh-treated mice (timecourse). & denotes *p*<0.1 vs. WT overall as indicated.

### Brain region SUV comparison at steady-state levels

Given the evident SUV difference in expression overtime in each brain region, the last 5 minutes of each scan were analyzed for comparative steady-state levels. In the dorsal striatum, a main effect of gene [F(1,11)=7.8, *p*<0.05; Fig. 4A] revealed elevated SUV in gp120 vs. WT littermate mice, while a LPS by gene interaction was observed [F(1,11)=5.1, *p*<0.05], although *post hoc* analyses revealed no effect of LPS in WT mice (F<1, ns), or gp120 Tg mice [F(1,5)=3.5, *p*=0.118]. In the ventral striatum, a trend increase in SUV was seen in gp120 Tg vs. WT littermate mice [F(1,11)=4.0, *p*=0.070; Fig. 4B]. While again no effect of LPS on WT mice were observed (F<1, ns), LPS increased ventral striatal SUV levels in gp120 Tg mice [F(1,5)=13.8, *p*<0.05]. In the hypothalamus, gp120 Tg mice exhibited elevated SUV levels vs. WT littermate mice [F(1,11)=6.7, *p*<0.05; Fig. 4C], and a gene by LPS interaction was also observed [F(1,11)=6.8, *p*<0.05]. As with the ventral striatum, this interaction was driven by LPS-induced increase in SUV in gp120 Tg mice [F(1,5)=5.3, *p*=0.070], an effect not seen in WT mice (F<1, ns). In the VTA, gp120 Tg mice exhibited elevated SUV levels overall vs. WT littermate mice [F(1,11)=6.3, *p*<0.05; Fig. 4D], with a gene by LPS interaction also observed [F(1,11)=6.6, *p*<0.05]. As before *post hoc* analyses revealed that no LPS-induced increase was seen in WT mice (F<1, ns), but a trend increase was seen in gp120 Tg mice [F(1,5)=4.4, *p*=0.091]. Finally, gp120 Tg mice continued to exhibit elevated SUV levels in the PFC vs. WT littermate mice [F(1,11)=6.2, *p*<0.05; Fig. 4E], but unlike other regions no gene by LPS interaction was observed (F<1.8, ns).

**Figure. 4.**
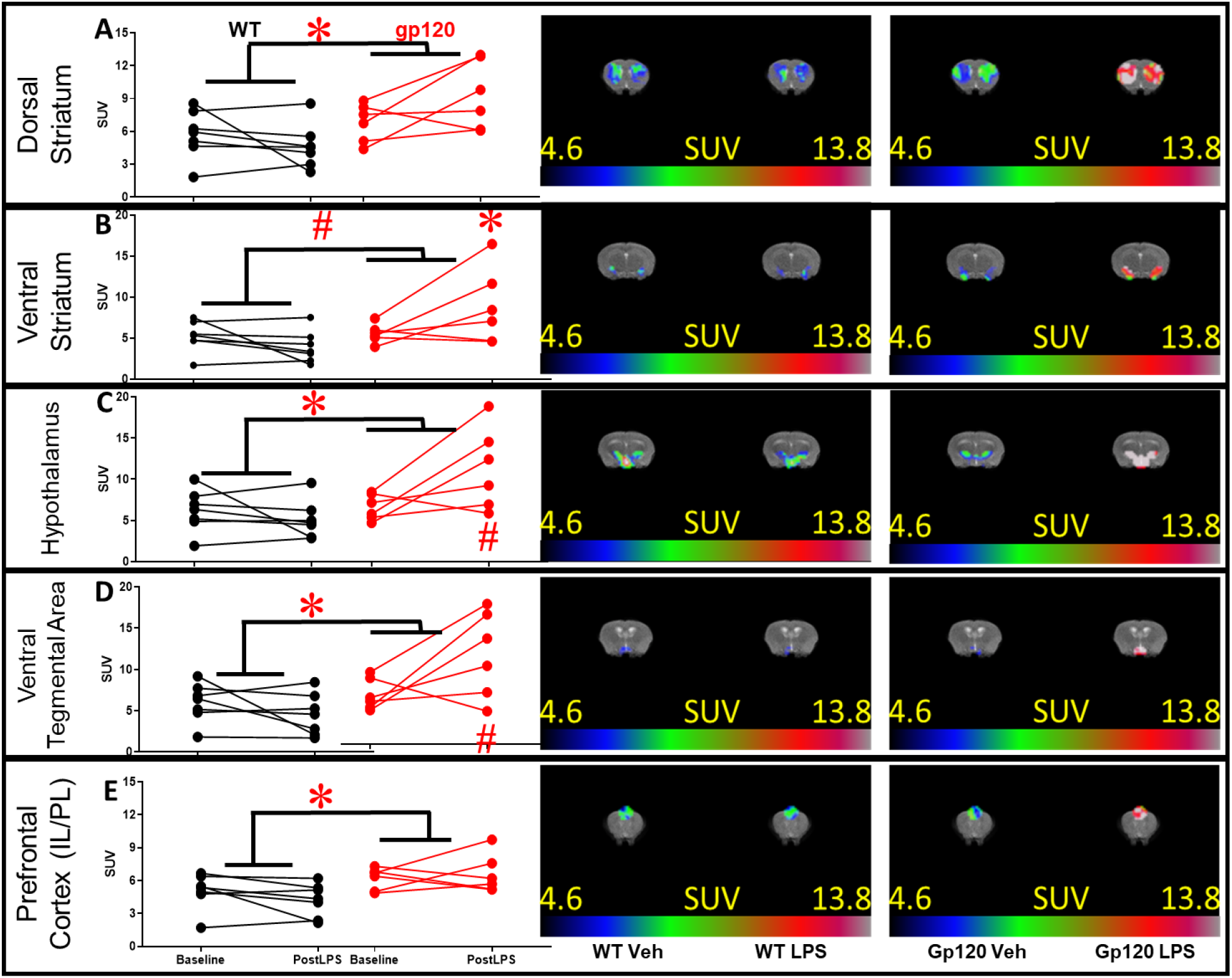
Mean micro-PET outcome of [(18)F]FEPPA studies for the final 5 min in each brain region. Gp120 Tg mice exhibited elevated overall SUV vs. WT littermates in the dorsal striatum (**A**), ventral striatum (**B**), hypothalamus (**C**), ventral tegmental area (**D**), and prefrontal cortices (PFC; **E**). These main effects were largely driven by LPS-induced increased SUV in gp120 Tg mice, not seen in WT mice across brain regions, although not in the PFC. Data presented as individual data points after vehicle (Baseline) and after LPS treatment. Images of SUV-activity from each brain region is provided * = p<0.05 as indicated, or vs. at baseline within genotype. # = p<0.01 as indicated, or vs. at baseline within genotype.

## Discussion

Here, we demonstrate the utility of *in vivo* [(18)F]FEPPA microPET for longitudinally assessing neuroinflammation in mice. We report that the neuroinflammatory response to an immune challenge (LPS) is heightened in gp120 Tg mice (an animal model relevant to HIV) compared to their WT littermates as measured by the microglial activation marker TSPO. Importantly, LPS decreased weights in both the WT and Tg mice, but the LPS-induced neuroinflammation was only observed in the gp120 Tg mice. This neuroinflammatory effect observed in gp120 Tg mice is similar to the elevation in PET-derived TSPO binding observed in PLWH, even when these individuals are treated with ART and have viral loads close to zero ^39, 40^. These data support the use of gp120 Tg mice to delineate the neural mechanisms driving the HIV-relevant gp120-induced sensitivity to neuroinflammation and the development of potential treatments.

LPS exerts numerous effects beyond its inflammatory response, including a general decrease in health ^41^. The efficacy of the LPS treatment was confirmed in both genotypes, as both WT and gp120 Tg mice lost equal amounts of weight following LPS administration (Figure 2B). Thus, LPS affected the health of both genotypes equally, consistent with previous studies^36^. The selective LPS-induced increase in TSPO levels in gp120 Tg mice were, therefore, likely a result of the interactive effects of LPS on neuroinflammation in these mice, not simply a reduced sensitivity to this treatment in the WT littermates.

*Ex vivo* histology can produce more sensitive findings than *in vivo* PET imaging. For example, elevated brain lysate levels of the proinflammatory cytokines IFN-γ, TNF-α, and IL-1β were observed in HIV-1 tg rats ^42^. The *in vivo* PET radiotracer [(18)F]-DPA714, however, failed to reveal differences in neuroinflammation at 3, 9, or 16 months of age ^43^ in these rats, although *ex vivo* Iba1 levels also did not differ in the latter study. In other disease models, such as, the APP/PS1-21 mouse model of Alzheimer’s disease, the radiotracer of TSPO using [18F]F-DPA714 revealed elevated microglial activation *ex vivo* at 4.5 mo, while *in vivo* PET evidence was not observed until 9 mo of age ^44^. Previous studies have revealed heightened neuroinflammation in other animal models of Alzheimer’s disease ^44-51^, ischemic stroke (^52^; ^53^), multiple sclerosis ^54^, and head injury ^55^. Given that neuroinflammatory differences in *ex vivo* tissue have been noted between gp120 Tg and WT littermate mice at baseline ^56^ (an effect somewhat observed here, but more pronounced after LPS treatment), these and our current data support the lower sensitivity of *in vivo* PET imaging to reveal neuroinflammatory differences. Further, greater differences in neuroinflammation between gp120 and WT littermates was also observed with increased age, as have cognitive deficits^56^. This study remains the first to reveal an *in vivo* heightened sensitivity of a mouse model of HIV to external infections. Future research determining the neuroinflammatory profile of these mice should include long-term aging and cognitive/behavioral testing with relevance to HIV-associated neurological disorder.

Unlike *ex vivo* assessment of neuroinflammation, *in vivo* measurement of TSPO using PET imaging techniques described here enable the longitudinal assessment of neuroinflammation potentially coinciding with behavioral changes. For example, cognitive performance was correlated with neuroinflammation in the PS2APP animal model of Alzheimer’s disease, wherein TSPO was correlated with spared cognitive performance as the animals aged ^57^. Furthermore, studies can be conducted in response to treatments, that may alter the neuroinflammatory profile e.g., LM11A-31 ^49^. For example, it remains possible that this heightened neuroinflammatory sensitivity of gp120 Tg mice to LPS is a result of impacted blood brain barrier permeability in these mice, relative to their WT littermates. Indeed, the BBB permeability is increased in gp120 Tg mice due to circulating gp120 protein ^58^, and may explain differences in radiotracer clearance. In another model of HIV, EcoHIV mice, BBB permeability is compromised and treatment with the antiplatelet drug eptifibatide restored this function ^59^.

Some evidence exists that gp120 interactions with alpha7 nicotinic acetylcholine receptors may contribute to this permeability ^60^. Alternatively, there remains evidence that the neuronal cell loss and neuroinflammation seen *ex vivo* in gp120 Tg mice could be blocked after administration of the immunomodulatory drug FK506 ^61^. Additionally, cognitive performance could be connected to other pathogenic factors of HIV. For example, HAND is also characterized by a reduction in synaptic density, loss of neurons, and astrocyte dysfunction, all possible contributing factors to neurocognitive decline ^62^. Thus, future studies measuring neuroinflammation in this mouse model of HIV could include FK506, antiplatelet, or nicotinic treatments to determine whether such hypersensitivity to LPS could be blocked or was a result of elevated BBB permeability, vs. a hypersensitivity to infection.

Limitations of this study include the use of a single 24 h timepoint, based in-part on previous literature. The lack of neuroinflammatory response in WT mice may have been a result of a different timescale driving neuroinflammation in WT vs. gp120 Tg mice. Another study revealed LPS-induced TSPO response 6 h after treatment in both male and female young (2 mo) and old (22 mo) mice, albeit from *ex vivo* tissue. Importantly, this study revealed no change in neuroinflammation or increased pro-inflammatory cytokine expression in young adult mice treated with LPS, but elevated neuroinflammation was observed in older mice (22 mo; ^63^). These findings support the present study of limited effects of LPS on neuroinflammation in younger mice. They highlight an additional limitation, however, in that only males were used in the current study, while Murtaj et al, 2019 observed a hypersensitivity with age in female mice. Likewise, HIV affects both males and females, inducing neuroinflammation irrespective of gender. Hence, future studies will endeavor to include both sexes as well as multiple ages. The validity of TSPO as a marker of inflammation has been under debate as it is not specific only to activated microglia, the main source of cytokines, but also to astrocytes and blood cells ^64^. However, TSPO is still consistently elevated in many disorders relating to neuroinflammation.

Furthermore, TSPO binding does decrease in patients post treatment showing that it can be used as a measure of clinical progress. In addition, this study was limited by the lack of blood analyses, which would have allowed for the calculation of total volume of distribution as an outcome measure, thereby controlling for individual differences in radiotracer metabolism and binding to vascular endothelium and plasma protein ^65-67^. Blood analyses were not performed however, as this is not feasible for small animal PET studies. Future studies will include immunohistochemistry studies to confirm presence of activated microglia in brain tissue as another confirmation of TSPO expression.

In summary, the present findings confirm that – consistent with human PET studies of PLWH – the gp120 Tg mouse model of HIV exhibits neuroinflammation, more specifically a heightened neuroinflammatory profile in response to an exogenous agent (LPS). This work confirms the sensitivity of microPET to detect such effects, as well as its utility for conducting longitudinal testing. Future studies will endeavor to incorporate cognitive testing relevant to HAND in addition to potential treatments that may remediate such neuroinflammatory and cognitive deficits.

